# Identification of a novel myosin antigen activating CD8^+^ T cells in C57BL/6 mice

**DOI:** 10.1101/2024.07.26.605251

**Authors:** Leon Richter, Michelle Bauer, Chiara Bernsen, Elena Vogel, Zabed Mahmud, Antoine-Emmanuel Saliba, Richard Schulz, Stefan Frantz, DiyaaElDin Ashour, Gustavo Campos Ramos

**Author notes:** **Corresponding authors** Prof. Dr. Gustavo Ramos, Immunocardiology Lab, Am Schwarzenberg 15, Haus A15, D-97078 Würzburg, Germany; Dr. DiyaaElDin Ashour, Immunocardiology Lab, Am Schwarzenberg 15, Haus A15, D-97078 Würzburg, Germany.

## Abstract

Heart-specific T cells play important roles in myocardial diseases, including myocarditis, myocardial infarction, and heart failure. However, only few cardiac antigens and their respective specific T cell receptors (TCRs) have been validated so far. Therefore, we sought to identify new cardiac T cell epitopes in C57BL/6 mice and to characterize the specific TCRs. Antigen mapping of myosin heavy chain alpha (MYHCA) led to the identification of a 9-mer epitope (MYHCA_724-732_) recognized by CD8^+^ T cells. Single-cell TCR sequencing (scTCR-seq) analysis of MYHCA-stimulated CD8^+^ T cells enabled the identification of distinct TCRs enriched on specific T cells. Selected TCRs inferred from scTCR-seq data were then expressed on reporter cell lines and functionally validated. Our study expands the toolkit required to study heart-directed immune responses by revealing a new cardiac antigen and specific TCRs. These tools will enable *in vivo* characterization of myosin-specific CD8^+^ T cells in murine models of myocardial diseases.

## Introduction

Antigen-specific T cells can play crucial roles in various myocardial diseases, including myocardial infarction,^1–4^ pressure-overload-induced heart failure,^5–7^ experimental autoimmune myocarditis and immune-checkpoint-inhibitor-induced myocarditis.^8,9^ T cells are activated upon antigen recognition through their somatically-rearranged T cell receptors (TCR). Therefore, identifying relevant cardiac antigens and developing tools to track heart-specific T cells are crucial for gaining a mechanistic understanding of the immunological basis of myocardial diseases.

Previous independent studies have identified the cardiac isoform of myosin heavy chain alpha (MYHCA) as a dominant cardiac antigen in various myocardial disease models.^4,8,10–18^ MYHCA (product of the *Myh6* gene) is highly expressed in adult murine ventricles and shares approximately 93% homology with the myosin heavy chain beta isoform (MYHCB, product of the *Myh7* gene), which is predominantly found in murine skeletal muscle tissue.^19^ Besides being an abundant cardiac protein, MYHCA also holds immunological significance. Unlike most tissue-specific proteins, MYHCA is not expressed in thymic epithelial cells.^18^ This lack of expression prevents MYHCA-specific T cells from undergoing thymic negative selection, thus, becoming more easily available in the periphery.

Most of the MYHCA antigens identified thus far have been described in BALB/c or A/J mouse strains, which are susceptible to developing autoimmune myocarditis in response to infection with cardiotropic viruses or to immunization with myosin.^4,8,10–14^ Immunization with MYHCA_614-643_ peptide in complete Freund’s adjuvant has been shown to trigger myocarditis in BALB/c mice.^12^ Building on these observations, Ludewig’s team developed transgenic mice (on BALB/c genetic background) in which CD4^+^ T cells are engineered to express a TCR specific for MYHCA_614-629_ presented on major histocompatibility complex (MHC) class II, resulting in the development of spontaneous autoimmune myocarditis.^8^ In A/J mice, the MHC-II restricted peptide fragment MYHCA_334-352_ has been shown to stimulate CD4^+^ T cells, leading to autoimmune myocarditis.^11,15^

In sharp contrast to A/J and BALB/c mice, C57BL/6 mice do not develop autoimmune myocarditis in response to immunization with myosin or to infection with cardiotropic viruses.^10^ The differences in disease susceptibility observed in these strains were mapped to differences in their MHC haplotypes; hence, to restrictions in antigen presentation.^20^ However, the recent discovery of a new form of fulminant myocarditis induced by immune checkpoint inhibitors has provided new insights into the pathophysiology of heart-directed immune responses.^9,21^ In particular, the recent discovery that *Pdcd1^−/−^ Ctla4^+/−^* mice on the C57BL/6 genetic background develop lethal CD8^+^ T cell-dependent myocarditis has reignited the interest in cardiac epitope mapping on this genetic background.^17,22^

The C57BL/6 mouse strain is the most common animal model used in cardiovascular and cardio-immunology basic research, with numerous transgenic substrains available for *in vivo* mechanistic studies. Therefore, we sought to identify new cardiac epitopes and generate new transgenic TCR tools for modeling disease in C57BL/6 mice. After combining antigen mapping and single-cell TCR sequencing (scTCR-seq) of MYHCA-stimulated T cells, we identified a novel cardiac epitope (MYHCA_724-732_) recognized by CD8^+^ T cells and validated the corresponding specific TCRs. These findings provide rationale and new tools for the *in vivo* examination of heart-specific T cell responses in a broad range of cardiac pathologies.

## Results

### Identification of MYHCA_724-732_ as a myosin epitope in C57BL/6 mice

To identify novel cardiac antigens in C57BL/6 mice, we initially queried the full length MYHCA amino acid sequence on the Immune Epitope Database (IEDB) to identify specific peptides predicted to bind to MHC-II molecules. The search parameters were adjusted to predict the binding of 15-mer peptides to murine I-A^b^, the MHC-II haplotype present in C57BL/6 mice.^4,23^ Our decision to initially focus on MHC-II restriction first stemmed from our previous studies on myosin-specific CD4^+^ T helper cells in BALB/c mice.^4,14^ Based on the *in silico* prediction results, we selected the 10 MYHCA peptides with the highest predicted binding probability (Figure 1 A, Table S1 A) to be synthesized and tested in antigen stimulation assays *in vitro* (Figure 1 B).

**Figure 1:**
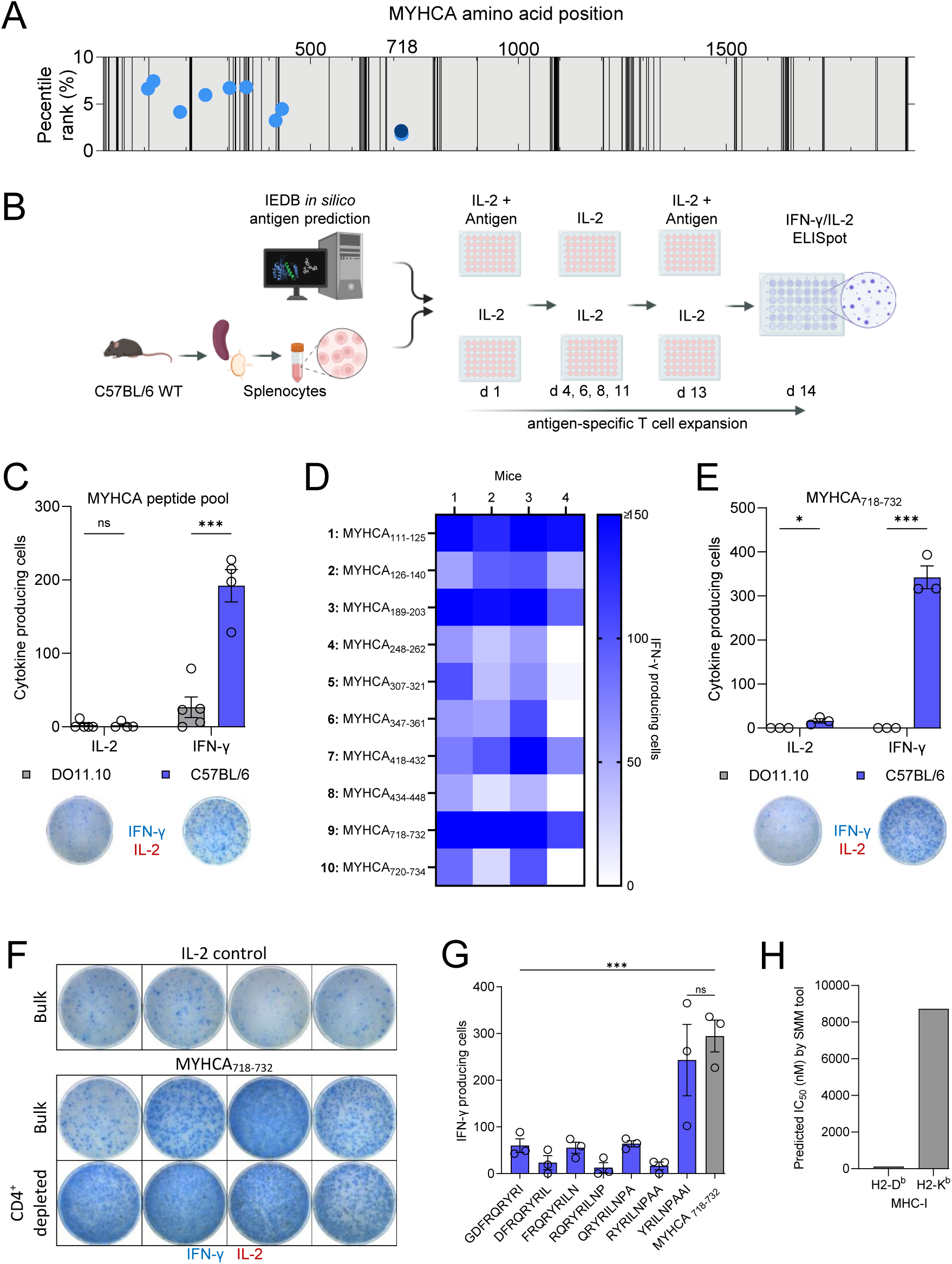
Identification of MYHCA_724-732_ as myosin epitope in C57BL/6 mice. **A:** MYHCA derived peptide fragments with the highest *in silico*-predicted binding to MHC I-A^b^ (blue dots; MYHCA_718-732_: dark blue). The grey areas indicate homologous regions between MYHCA (cardiac isoform) and MYHCB (skeletal muscle isoform), whereas the black lines indicate non-homologous regions. The amino acid positions are plotted on the X-axis, while the percentile rank for the peptide binding to I-A^b^ is plotted on the Y-axis. **B:** Based on the MHC-II binding predictions, the top 10 MYHCA peptides were synthesized and tested *in vitro* using C57BL/6 splenocyte preparations. IL-2 and IFN-γ production upon 14-day antigenic stimulation was assessed by ELISpot assay. **C:** Quantification of IL-2 and IFN-γ producing cells upon stimulation with a pool of 10 MYHCA peptides. Splenocytes prepared from DO11.10 mice, which express a transgenic TCR specific for an irrelevant antigen (OVA_257-264_), were used as control. ELISpot pictures are representative for the biological replicates; *n* = 5 DO11.10 mice and *n* = 4 C57BL/6 mice; data acquired from two independent experiments. **D:** Splenocytes were challenged with individual MYHCA peptides, and the IFN-γ production was measured by ELISpot assay; *n* = 4 C57BL/6 mice. **E**. Splenocytes from C57BL/6 and DO11.10 mice were stimulated using the MYHCA_718-732_-fragment and the IL-2 and IFN-γ production were measured by ELISpot. The ELISpot pictures are representative for the biological replicates; *n* = 3 DO11.10 and C57BL/6 mice. **F:** Cytokine production was also analyzed on CD4^+^ T cell-depleted splenocytes, as compared to bulk splenocyte preparations; *n* = 4 C57BL/6 mice for bulk and CD4^+^ T cell-depleted. **G:** IFN-γ production by CD4^+^ T cell-depleted splenocytes stimulated with MYHCA_718-732_ or each of its seven 9-mer peptide-fragments; *n* = 3 C57BL/6 mice. **H:** Binding prediction of YRILNPAAI-peptide to C57BL/6 MHC-I haplotypes using IEDB analysis resource SMM tool.^49^ A lower IC_50_ value indicates higher binding affinity of the peptide to the respective MHC-I molecule. The graphs show the individual distribution of *n* datapoints alongside with mean ± S.E.M. Statistical analysis: unpaired t test with correction for multiple comparisons by Holm-Sidak method (for C and E); One-way ANOVA with Dunnett’s multiple comparisons test (for G). *, p < 0.05; **, p < 0.01; ***, p < 0.001.

Next, splenocytes isolated from naïve C57BL/6 mice were cultured for 14 days in the presence of IL-2 and a peptide pool containing all 10 MYHCA candidate epitopes identified *in silico*. As control, splenocytes were cultured in the presence of IL-2 only. This long-term *in vitro* stimulation protocol allows for clonal expansion of specific T cells initially present at low frequencies, which could then be detected based on interferon-γ (IFN-γ) and interleukin-2 (IL-2) production upon antigen recall (Figure 1 B). As shown in Figure 1 C, long-term *in vitro* stimulation with a MYHCA peptide pool gave rise to significantly elevated numbers of IFN-γ-producing cells in splenocytes prepared from C57BL/6 compared to DO11.10 splenocytes, which express a transgenic TCR against OVA_323-339_ peptide presented on I-A^d^ (BALB/c background). These findings indicate an antigenic capacity of MYHCA peptide(s) present within the tested pool. Further analyses with peptides tested individually revealed four potentially interesting MYHCA peptides accounting for the T cell stimulation in this system, including peptides 1 (MYHCA_111-125_), 3 (MYHCA_189-203_), 7 (MYHCA_418-432_) and 9 (MYHCA_718-732_) (Figure 1 D). Interestingly, the peptides 3 (MYHCA_189-203_) and 7 (MYHCA_418-432_) contained 8-mer sequences that overlap with MHC class-I-restricted epitopes recently identified by Axelrod, et al, in the context of immune-checkpoint-inhibitor-induced myocarditis.^17^ Based on these observations, we decided to focus on the peptide MYHCA_718-732_, which showed the strongest and most consistent antigenicity *in vitro*. Notably, three-dimensional structure prediction of the motor domain of myosin S1 fragment maps MYHCA_718-732_ to an exposed position that is likely readily accessible to proteases. Furthermore, the structural context of the MYHCA_718-732_-epitope and its proximity to the actin-binding site MYHCA_759-773_ is highlighted (Figure S1).

Additional experiments confirmed C57BL/6 but not DO11.10 splenocytes were able to respond to MYHCA_718-732_ stimulation, indicating a role of MHC restriction in these observations (Figure 1 E). Next, we repeated the experiments on CD4^+^ T cell depleted or bulk splenocyte populations to further characterize the MHC restriction in this model. Contrary to our expectations, and despite the tested epitopes being selected based on predicted MHC-II binding, ELISpot analyses revealed that the CD8^+^ T cell compartment accounted for MYHCA_718-732_-induced IFN-γ production (Figure 1 F). Because MYHCA_718-732_ comprises a 15-mer amino acid sequence, and considering that antigen presentation to CD8^+^ T cells in the context of MHC-I normally harbors shorter peptides,^24–26^ we generated a peptide library with all possible 9-mer combinations within this sequence (Table S1 B). Additional *in vitro* stimulation experiments followed by ELISpot analysis revealed that IFN-γ response to the shorter peptide MYHCA_724-732_ (YRILNPAAI) was comparable to that observed for MYHCA_718-732_ (Figure 1 G). Additionally, analysis on the IEDB Analysis Resource predicted that MYHCA_724-732_ can bind with high affinity to the class-I MHC of H2-D^b^ (IC_50_ = 117.9 nM) (Figure 1 H). In conclusion, we identified MYHCA_724-732_ as a novel MHC-I-restricted antigen able to stimulate CD8^+^ T cells obtained from C57BL/6 mice.

### MYHCA-stimulation induces T cell activation and expansion

After having identified a new MYHCA epitope, we sought to characterize the specific TCRs recognizing this antigen. Therefore, we isolated splenocytes from C57BL/6 mice expressing yellow fluorescent protein (YFP) reporter under the control of *Ifng* promotor (*Ifng*^YFP^).^27^ To focus on specific TCRs expressed on CD8^+^ T cells, we depleted CD4^+^ T cells using magnetic activated cell sorting prior to antigenic stimulation. After 14-day stimulation with MYHCA_718-732_ peptide the TCRβ^+^CD8^+^CD44^high^IFN-γ-YFP^+^ cells were purified by fluorescence-activated cell sorting (FACS) and prepared for downstream single-cell RNA/TCR-sequencing (scRNA/TCR-seq). As controls, TCRβ^+^CD8^+^CD44^+^ cells from IL-2-stimulated cells were also purified (Figure 2 A). Since IL-2 treatment did not induce IFN-γ expression, it was not feasible to enrich for IFN-γ-YFP^+^ under this condition.

**Figure 2:**
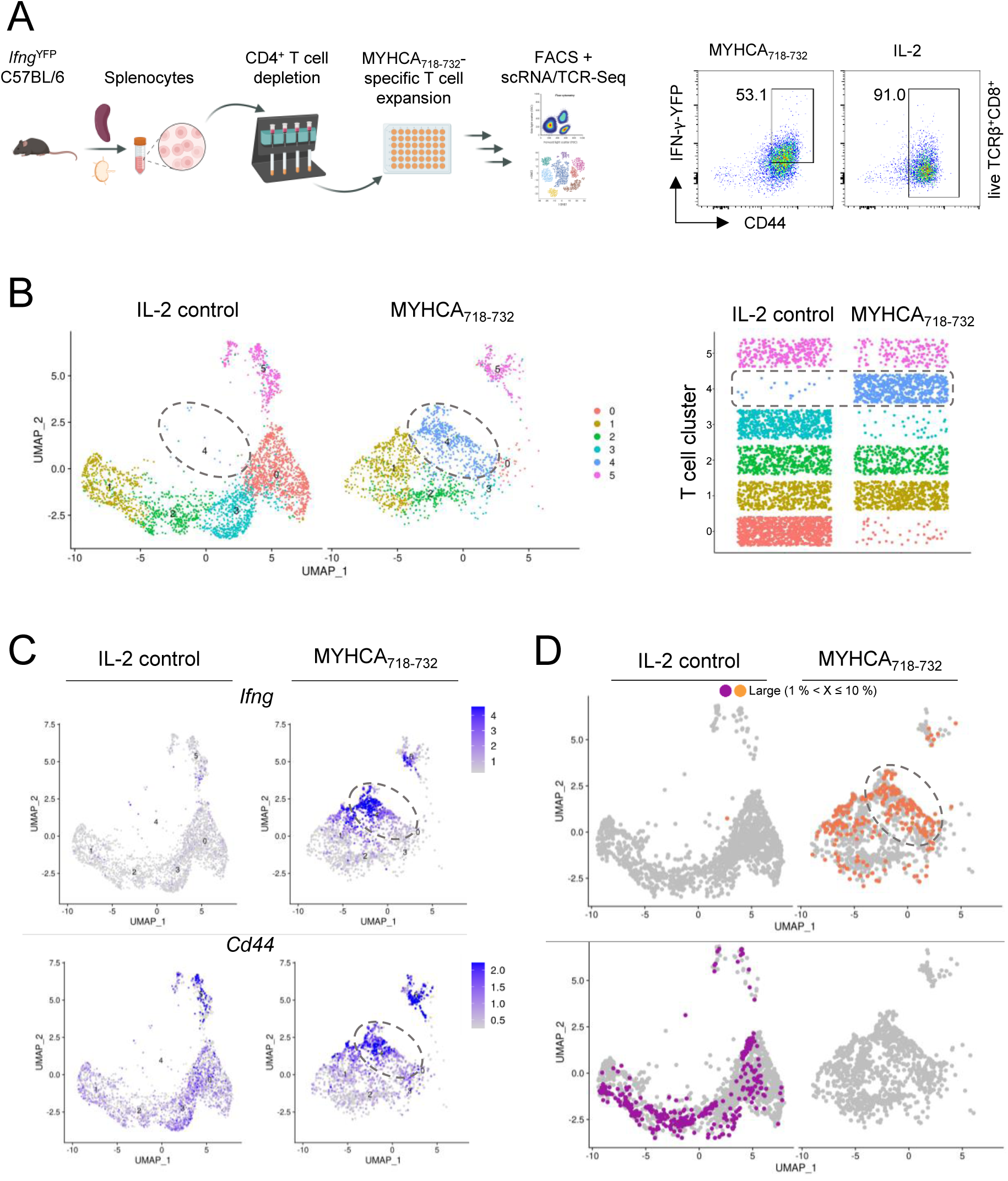
MYHCA-stimulation induces CD8^+^ T cell clonal expansions. **A:** Workflow for *in vitro* expansion of MYHCA_718-732_-specific CD8^+^ T cells from C57BL/6 I*fng*^YFP^ reporter mice and sample preparation for scRNA/TCR-seq. Spleen and lymph nodes from *Ifng*^YFP^ C57BL/6J mice were processed to single-cell suspensions and the CD4^+^ T cells were depleted by magnetic activated cell sorting. After 14d-*in vitro* stimulation with MYHCA_718-732_, or IL-2 control, cells were purified based on their expression of activation markers CD44 and YFP, and then prepared for scRNA/TCR-seq. Representative FACS gating plots are shown. **B:** UMAP representation of 4,806 single cell transcriptomes showing clusters identified based on differentially expressed gene signatures (left) and Jitter-plot showing the distribution of cells for each cluster (y-axis) and condition (x-axis) (right). Cell cluster 4 (highlighted) distinguished cells from MYHCA_718-732_-stimulated vs IL-2 conditions. **C:** Expression levels projected on the UMAP as displayed in (B) of activation markers *Cd44* and *Ifng* in control and MYHCA_718-732_-stimulated cells. **D:** Identification of highly expanded clones (representing unique TCRs shared by > 1% of total cells) in control and MYHCA_718-732_-stimulation conditions projected on the UMAP. T cell clones marked in orange were expanded upon MYHCA_718-732_-stimulation, whereas the T cell clones shown in violet were found expanded in IL-2 control conditions.

The TCRβ^+^CD8^+^CD44^high^IFN-γ-YFP^+/-^ T cells purified from IL-2 (control) and MYHCA-stimulated conditions were multiplexed using barcoded hashtag antibodies (anti-CD45/MHC-I TotalSeq-C) and sequenced as a single library. A total of 6,014 purified T cells were sequenced. After conducting a quality check and excluding cells with double hashtag signals or no hashtag detection and applying quality control filters for RNA content and gene count, 4,806 cells remained for downstream analysis (Figure S2 A). Clustering analysis based on the gene expression profile of all cells revealed 6 distinct T cell clusters (Figure 2 B, Figure S2 B). The distribution of cells differed between the studied conditions, with cluster 4 being almost exclusively present in the MYHCA_718-732_-stimulated condition (Figure 2 B). A more detailed analyses of the CD8^+^ T cells found in cluster 4 revealed that they were enriched for transcripts associated with TCR activation (e.g. *Cd44*) and effector cytokines (e.g. *Ifng*) (Figure 2 C). In addition, differential gene expression analysis revealed that several chemokines (e.g. *Ccl1*, *Ccl3*, *Ccl4*, and *Xcl1*), were among the top expressed gene transcripts in cells from cluster 4. *Crtam*, a gene expressed in activated CD8^+^ T lymphocytes also adds to this list (Figure S2 B).^28^ These findings indicate that cluster 4 might comprise a population of antigen specific CD8^+^ T cells activated by MYHCA_718-732_-stimulation.

TCR repertoire analyses further confirmed that cluster 4 comprises a subset of highly expanded CD8^+^ T cell clones exclusively found upon MYHCA stimulation. IL-2 stimulated control cells also showed some clonal expansion, though with a different distribution. Importantly, we did not observe overlaps between the TCR repertoires of MYHCA_718-732_ vs IL-2 treated cells (Figure 2 D). Based on the transcriptional profile and clonal expansion levels of MYHCA-stimulated cells, we concluded that the top expanded TCRs found in cluster 4 might confer specificity to the MYHCA antigen herein identified. Therefore, our subsequent analysis of specific TCR motifs focused on this cell population.

### TCR repertoire analysis suggests MYHCA-specific T cell expansion

The antigen-specificity of a given TCR is primarily defined by its complementarity determining region 3 (CDR3), which contacts the peptides presented on MHC.^29^ Therefore, we focused our analyses on the TCRβ CDR3 sequences. First, we computed the Levenshtein distances among the TCRβ CDR3 sequences of the top 20 most expanded clones found in the control and stimulated conditions. The Levenshtein distance between two TCRs is defined based on the number of amino acid substitutions, insertions, or deletions observed between their CDR3 sequences (where each difference increments the Levenshtein distance by 1 for every amino acid mismatch).^30,31^ TCR sequences found in MYHCA_718-732_-stimulated, but not IL-2-treated, CD8^+^ T cells showed a great degree of TCR convergence, suggesting that they may recognize a conserved epitope (Figure 3 A).

**Figure 3:**
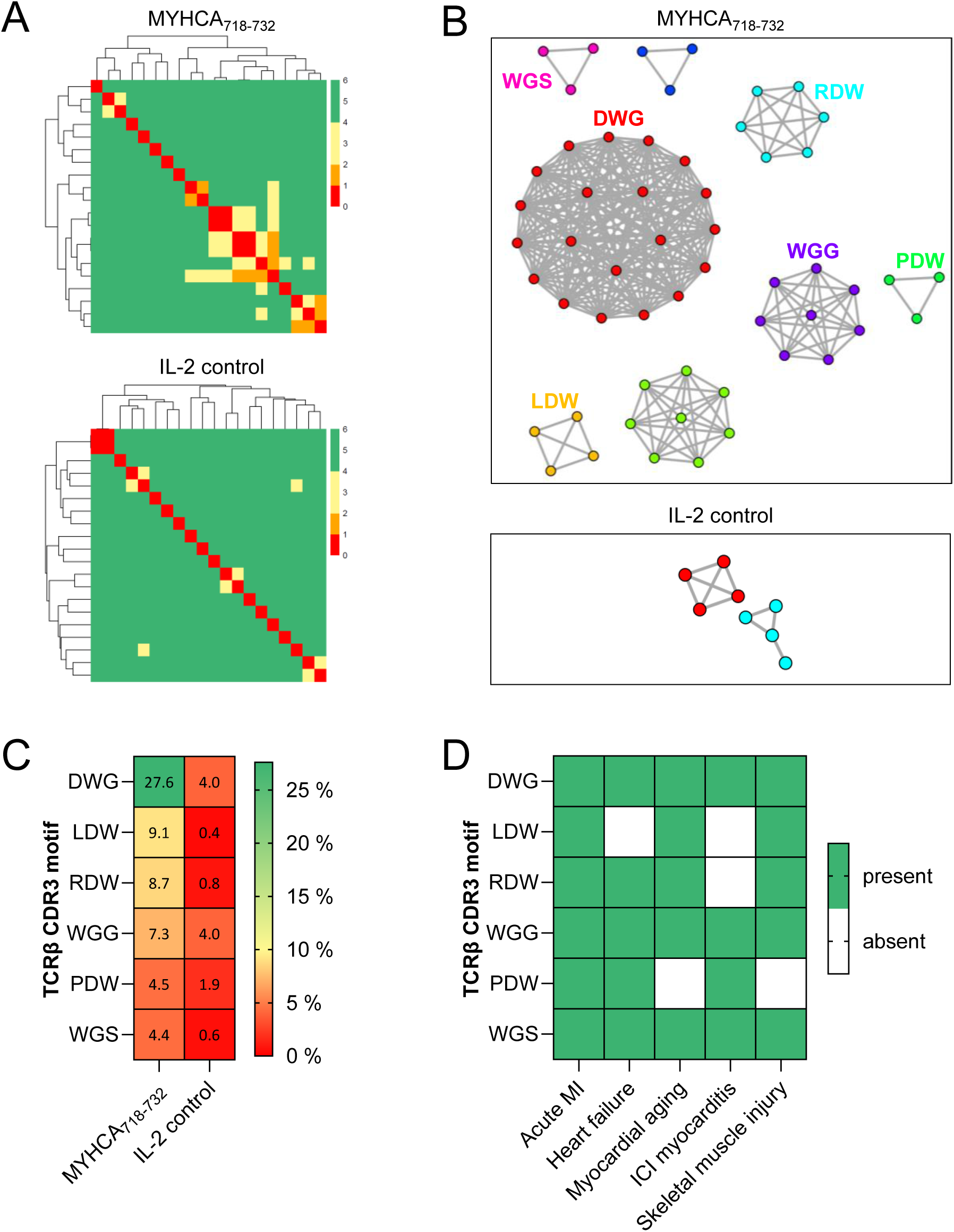
TCR repertoire analysis suggests MYHCA-specific T cell expansion. **A:** Levenshtein distance analysis comparing the TCRβ CDR3 amino acid sequences of the top 20 most expanded clones present in IL-2 control or MYHCA_718-732_-stimulation conditions. Smaller Levenshtein distances indicate more similar TCR sequences. The color legend indicates the number of amino acid substitutions between two TCRs. **B:** GLIPH2 motif-enrichment-network analysis of the top expanded TCRβ CDR3 sequences in MYHCA_718-732_-stimulated and IL-2 control datasets. The networks connect unique TCRβ CDR3 sequences that share a common core motif. The CDR3 motifs for selected networks are given in matching colors. A two-fold enrichment against motifs generated from a control dataset of 78,116 TCRβ CDR3 sequences present in healthy mice was used as cutoff for cluster generation. **C:** CDR3 motifs from the MYHCA_718-732_-GLIPH2 analysis were selected and checked for their occurrence in TCRβ CDR3 sequences from each MYHCA_718-732_-stimulated or IL-2 control cell. The heatmap colors indicates the relative frequencies of TCR motifs in each experimental group. **D:** Meta-analysis of TCR repertoire datasets available from previous studies. The top expanded CDR3 sequences found in different models of myocardial disease and skeletal muscle injury datasets were screened for selected MYHCA_718-732_ CDR-3 motifs. The heat map indicates the presence or absence of selected CDR3 motifs in the various disease conditions. MI, myocardial infarction; ICI, immune-checkpoint-inhibitor.

Next, to identify common TCRβ CDR3 motifs potentially conferring MYHCA-specificity, we analyzed the TCR repertoire of MYHCA-stimulated cells based on Grouping of Lymphocyte Interactions by Paratope Hotspots (GLIPH2) motif-enrichment-network analysis.^32,33^ The GLIPH2 algorithm searches and groups TCRβ CDR3 sequences based on conserved motifs within the core-region of the CDR3 that has the highest contact probability with the antigen docked on the MHC molecule.^32^ The first and last 4 boundary amino acids of each CDR3 were excluded from the analysis as they mostly represent germline derived sequences, and 3 amino acid core motifs were used for network building. The top expanded TCRβ CDR3s from both MYHCA_724-732_- or IL-2 stimulation were compared against motifs enriched from a reference dataset comprising 78,116 control TCRs obtained from healthy C57BL/6 mice, to control for common CDR3 motifs.^33^ This approach allowed us to identify distinct TCR motifs potentially associated with myosin recognition (Figure 3 B). In the MYHCA-stimulated cell subset, almost 30% of cells carried a prominent ‘DWG’ motif in their core TCRβ CDR3 region. Several variations around this TCR core motif (i.e. ?DWG?) were also found expanded amongst MYHCA-stimulated, but not in control, conditions (Figure 3 C).

Based on the Levenshtein-distances and GLIPH2-network analyses we were able to identify conserved TCRβ CDR3 motifs enriched in CD8^+^ T cells stimulated with MYHCA_718-732_ antigen. To further assess the potential physiological relevance of these TCRs, we conducted a meta-analysis using several publicly available TCR repertoire datasets generated from myocardial and skeletal muscle injury models.^17,31,34–37^ Interestingly, the TCR motifs enriched in MYHCA_718-732_-stimulated CD8^+^ T cells were also observed among the top expanded clones in experimental models of acute myocardial infarction,^34^ pressure-overload-induced heart failure,^35^ immune-checkpoint-inhibitor-induced myocarditis,^17^ and skeletal muscle injury (Figure 3 D).^37^ These findings might hint at an *in vivo* relevance of the described MYHCA epitope and specific TCRs in various settings of muscle injury.

### Engineered MYHCA-TCR reporter T cells react to MYHCA_724-732_

Next, we proceeded to functionally validate the specificity of the TCRs identified from our scTCR-seq data by expressing candidate MYHCA-specific TCRs in engineered reporter T cell lines. We adapted a protocol established by the group of Maki Nakayama,^38,39^ and generated reporter T cell lines expressing selected TCRs, utilizing the thymoma cell line BW5147.G.1.4 as a starting point. However, due to the lack of surface CD3 and CD8 expression and the presence of endogenous TCR chains in these cells, we first needed to implement a series of modifications before expressing the TCRs of interest (Figure 4 A). First, we expressed the four mouse CD3 subunits using the CD3-2A_pMI-LO construct^38^ using murine MSCV retroviral vectors. Surface CD3 expression is required for TCR activation and the introduction of this construct also induced surface co-expression of the BW5147.G.1.4 endogenous TCR (Figure S3 A). Next, we employed CRISPR/Cas9 mediated excision of the TCR constant regions to prevent expression of the BW5147.G.1.4 endogenous TCRs (Figure S3 B). Afterwards, we retrovirally introduced the expression of mCD8 alpha and beta subunits, which are also required for recognition of antigens presented on MHC-I. The expressed mCD8 vector (8xNFAT-ZsG-mCD8^39^), additionally contained NFAT-driven ZsGreen-1 gene segments, serving as reporter for TCR stimulation (Figure S3 C).

**Figure 4:**
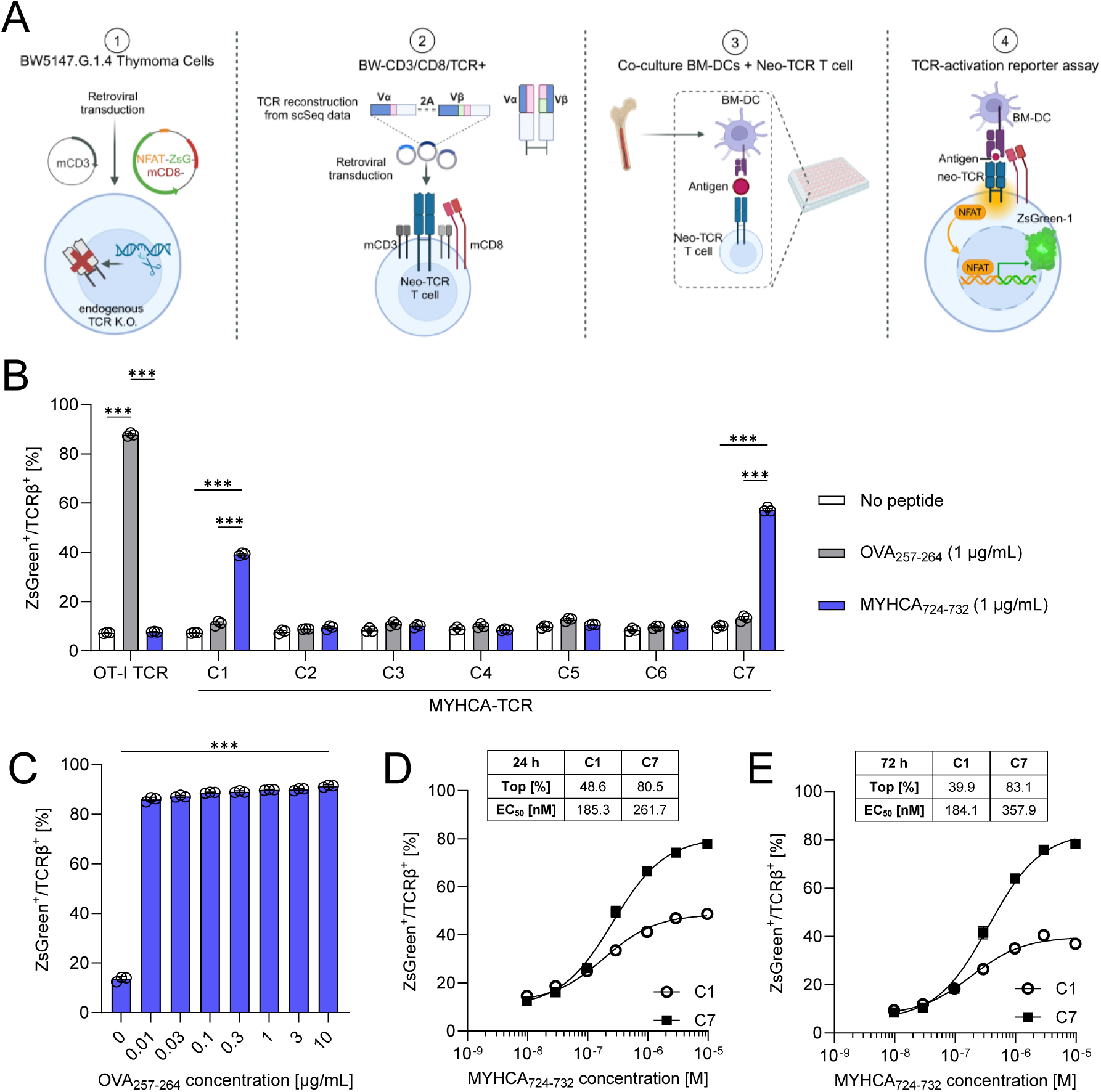
Engineered MYHCA-TCR reporter T cells react to MYHCA_724-732_. **A:** Generation of a T cell activation reporter assay and cloning of selected TCRs. (1) BW5147.G.1.4 thymoma cells served as starting point for TCR-engineered reporter T cell lines. The endogenous α and β TCR constant gene segments were subjected to CRISPR/Cas9 mediated excision and BW5147.G.1.4 thymoma cells were transduced with a retroviral vector containing the four chains of murine CD3^38^ and a vector containing NFAT-inducible ZsGreen-1 and murine CD8α and β subunits.^39^ (2) TCR α and β chains were reconstructed *in silico* from CDR3 sequences and V and J genes available from scTCR-seq experiments. TCR α and β chain genes were separated by a 2A sequence, cloned into retroviral vectors and used to transduce BW-CD3/CD8^+^ cells. (3) Reporter T cells were co-cultured with antigen-loaded bone marrow-derived dendritic cells (BMDCs). (4) Upon antigen recognition and TCR activation, NFAT transcription factor binds to the respective binding site and promotes ZsGreen-1 expression, which can be used as flow-cytometric readout for T cell activation. The OT-I TCR was expressed as control, since it confers specificity to a mapped ovalbumin epitope that is not present in mammalian hearts. **B:** MYHCA-TCR and OT-I-TCR reporter T cells were co-cultured for 24 h with BMDCs loaded with 1 μg/mL OVA_257-264_ or MYHCA_724-732_, respectively, or with unloaded BMDCs as control. ZsGreen signal was monitored by flow cytometry as readout for TCR-stimulation. **C:** OT-I-TCR clones were co-cultured for 24 h with BMDCs loaded with increasing concentrations of OVA_257-264_. **D-E:** MYHCA-TCR clones 1 and 7 were co-cultured for 24 h **(D)** or 72 h **(E)** with BMDCs loaded with increasing concentrations of MYHCA_724-732_. The graphs show the individual distribution of three technical replicates and/or mean ± S.E.M. Statistical analysis: Two-way ANOVA followed by Tukeýs multiple comparisons test (for B); One-way ANOVA with Dunnett’s multiple comparisons test (for C); nonlinear agonist vs. response fit for EC_50_ (for D and E). *, p < 0.05; **, p < 0.01; ***, p < 0.001.

Finally, to express candidate MYHCA-specific TCRs, we reconstructed the TCR α and β chains corresponding to the top seven expanded clones from our MYHCA-stimulated scTCR-Seq dataset (Table S2) using Stitchr^40^ and retrovirally transduced them into reporter 8xNFAT-ZsG-mCD8 + CD3-2A_pMI-LO BW5147.G.1.4 cells. To functionally interrogate specificity of all selected TCRs, syngeneic bone-marrow-derived dendritic cells loaded with the cognate antigen were used as antigen presenting cells and co-cultured with the various TCR_8xNFAT-ZsG-mCD8 + CD3-2A_pMI-LO BW5147.G.1.4 cell lines generated. The expression levels of the NFAT driven fluorescent reporter were monitored by flow cytometry as readout for TCR activation (Figure 4 A, Figure S3 D).

As proof of concept and to validate this in-house generated testing system, we expressed the well-characterized OT-I TCR that shows high affinity for the chicken ovalbumin peptide OVA_257-264_ (SIINFEKL).^41^ As shown in Figure 4 B-C and Figure S3 D, 8xNFAT-ZsG-mCD8 + CD3-2A_pMI-LO BW5147.G.1.4 cells expressing the OT-I TCR exhibited a dose-dependent increase in NFAT-ZsGreen expression upon stimulation with its cognate antigen SIINFEKL. Furthermore, no response was observed when these cells were co-cultured with unloaded antigen-presenting cells or stimulated with irrelevant peptide.

After having confirmed the functionality of the cell-reporter assay, we stimulated all 7 generated MYHCA-TCR reporter cell lines with MYHCA_724-732_ and then monitored their activation. As shown in Figure 4 B, MYHCA_724-732_-specific TCR stimulation could be confirmed for the clonotypes 1 (C1) and 7 (C7). In contrast, stimulation with an irrelevant peptide (OVA_257-264_) did not induce reporter signal equivalently, proving MYHCA-specific T cell activation for C1 and C7 (Figure 4 B). Next, to determine the half-maximal effective concentration of peptide (EC_50_), we performed a peptide titration experiment.^42,43^ In this context, we found that the MYHCA-specific clones C1 and C7 exhibited similar EC_50_ values (24 h: 185.3 nM and 261.7 nM; 72 h: 184.1 nM and 357.9 nM respectively), while differing in the maximal mounted responses (24 h: 48.6% and 80.5% activation; 72 h: 39.9% and 83.1% activation respectively) (Figure 4 D-E). The affinities observed for TCRs recognizing the autoantigen MYHCA were lower when compared to the OT-I TCR, which recognizes a non-self antigen. These observations confirm that our in-house generated reporter cell lines provide a suitable platform for expressing and validating TCRs inferred from scTCR-seq datasets. Eventually, this has led to the generation of two cell clones with confirmed specificity to the MHC-I-restricted MYHCA_724-732_ epitope.

## Discussion

In the present study we identified MYHCA_724-732_ (YRILNPAAI) as a cardiac antigen able to stimulate CD8^+^ T cells in C57BL/6 mice. Moreover, after performing scRNA/TCR-seq of CD8^+^ T cells stimulated *in vitro* with this MYHCA epitope, we identified conserved TCR motifs enriched in the antigen-specific T cells. Finally, by expressing selected TCRs on in-house generated reporter T cell lines, we were able to functionally validate the specificity of two MYHCA_724-732_ TCRs. The identification of novel cardiac antigens and the generation of new tools to track myosin-specific CD8^+^ T cells in C57BL/6 mice are significant steps in the field because few cardiac antigens have been identified in this mouse strain so far.^17^

BALB/c and A/J mice typically develop autoimmune myocarditis in response to immunization with myosin antigens in complete Freund’s adjuvant or infection with cardiotropic viruses.^10^ In BALB/c mice, MYHCA_614-629_ is now a well-established model cardiac antigen, with a transgenic TCR mouse model available,^8^ and validated MHC-II tetramers that allow mechanistic studies in different myocardial disease models.^14^ Likewise, MYHCA_338-348_ and MYHCA_334-352_ have been validated as model antigens in A/J mice.^13,15,16,44,45^ However, most studies in cardiology are performed in C57BL/6 mice, which do not develop myocarditis in response to myosin immunization or upon infection with cardiotropic viruses.^10^ These limitations have precluded the discovery of cardiac antigens and specific TCRs in this mouse strain until recently, thereby hindering developments in the field.

The discovery of a new form of myocarditis induced by immune checkpoint inhibition, including in *Pdcd1^−/−^Ctla4^+/−^* mice on a C57BL/6 background,^9,22^ has reignited interest in developing cardio-immunology tools to track specific T cells in C57BL/6 strains. Building on these new models of immune-checkpoint-inhibitor-induced myocarditis, Axelrod et al. identified two new MHC-I-restricted MYHCA epitopes in C57BL/6 mice (MYHCA_191-198_, MYHCA_418-425_).^17^ Our epitope mapping approach independently confirmed IFN-γ responses to similar MYHCA peptides (MYHCA_189-203_, MYHCA_418-432_) and revealed additional previously unrecognized MYHCA epitopes able to elicit even stronger responses *in vitro* (MYHCA_111-125_ and MYHCA_718-732_). However, it should be noted that our epitope mapping approach is far from exhaustive and additional MYHCA epitopes may still be identified in future studies.

Building on own previous studies focusing on myosin-specific CD4^+^ T cells in the context of myocardial infarction,^4,14^ we initially aimed at identifying MHC-II-restricted myosin antigens that could be presented to CD4^+^ T cells on I-A^b^. Surprisingly, despite testing 15-mer peptide fragments predicted to bind to MHC-II in our initial screening, our experiments using CD4^+^ T cell-depleted splenocytes revealed that the MYHCA epitope identified in this study is primarily recognized by CD8^+^ T cells. Thus, considering that antigens presented to CD8^+^ T cells on MHC-I are typically shorter than those presented on MHC-II,^24–26^ further refinements in our epitope mapping finally identified MYHCA_724-732_ as being the bona fide MHC-I-restricted antigen in our testing system. It is important to emphasize that the long-term *in vitro* stimulation approach used in this study can only reveal potential TCR reactivities that can be expanded from naïve repertoires. However, it does not provide definitive proof of the relevance of this epitope in specific disease conditions, which requires *in vivo* experimental validation.

Alongside with the identification of a new cardiac antigen, we employed scRNA/TCR-seq to characterize the TCR antigen-binding domains of myosin-specific CD8^+^ T cells. This approach enabled us to identify a defined subset of CD8^+^ T cells enriched for pro-inflammatory transcripts (e.g. *Ifng, Ccl1*, *Ccl3*, *Ccl4, Xcl1*) alongside with TCR-dependent activation markers (e.g. *Cd44*). Characterization of the TCR repertoire in this particular T cell subset revealed expansion of unique clones exhibiting some degree of TCR convergence (i.e. lower Levenshtein distances). Moreover, after comparing the CDR3 motifs of TCRs expanded upon MYHCA_724-732_-stimulation against a reference set comprising 78,116 control TCRs, we were able to identify distinct TCRβ CDR3 motifs potentially associated with myosin recognition. Interestingly a meta-analysis integrating various TCR repertoire datasets available in cardiology confirmed that the CDR3 motifs identified in our study were also expanded TCRs in various myocardial disease conditions.^17,31,34–36^ Given that multiple studies highlighted the importance of MYHCA epitopes to immune responses in cardiac pathologies,^4,8,10–18^ it is plausible to assume that MYHCA_724-732_-reactive T cells might also be physiologically relevant. However, future *in vivo* studies based on heart disease models are needed to confirm the relevance of the antigen and TCRs identified in our study.

Contrary to other MYHCA epitopes identified in BALB/c and A/J mice, which fall within non-homologous regions with MYHCB,^11–13,16^ the MYHCA_724-732_ peptide sequence identified in our study is identical to the MYHCB_722-730_ fragment. This is an important observation because MYHCA is not expressed by thymic epithelial cells whereas MYHCB is. This key difference may have important implications concerning the central immunological tolerance mechanisms to this antigen.^17,18^ Moreover, it is important to note that MYHCB is highly expressed in skeletal muscle,^46^ while MYHCA constitutes the main isoform in murine hearts.^47^ This could explain the putative expanded MYHCA_724-732_-specific CDR3 motifs were also observed in a dataset generated in a murine model of skeletal muscle injury.^37^

An important aspect of our study is the expression and functional validation of TCR motifs inferred from our scTCR-seq data using in-house generated T cell reporter cell lines. Using this “sequence and clone” approach, we confirmed and validated the MYHCA_724-732_-specificity of two TCR sequences. While differing in the maximal mounted response, both TCRs exhibited similar EC_50_ values and showed clear responses to antigenic stimulation at physiologically relevant concentrations (0.1-1 µg/ml). However, these myosin-specific TCRs exhibited higher EC_50_ values compared to the well-established OT-I TCR (specific for a non-self antigen). These differences in TCR affinity are not surprising, considering that MYHCA is a self-antigen.^48^ The functional validation of TCRs inferred from sequencing datasets underscores the value of modeling tools like GLIPH2. However, it is important to note that not all expressed TCRs exhibited the predicted binding properties. Thus, while current TCR modeling tools can provide valuable insights into TCR motif discovery, functional validation remains indispensable to confirm binding.

Taken together, we herein report the identification of MYHCA_724-732_ peptide fragment as MHC-I-restricted epitope able to trigger cytotoxic T cell responses in C57BL/6 mice. The identification of a novel class-I-restricted MYHCA antigen, together with the specific TCRs found to be expanded by antigenic stimulation, contributes to the development of a toolkit essential for cardio-immunology studies. These observations and tools herein presented pave the way for future studies aimed at characterizing heart-directed T cell responses in a variety of heart diseases, thus contributing to a comprehensive understanding of disease pathology.

## Acknowledgements

G.C.R. is supported by the German Research Foundation (Heisenberg Program, grant number 517001338) and the Interdisciplinary Center for Clinical Research Würzburg (Project S-511). A.E.S., S.F. and G.C.R. lead projects integrated in the Collaborative Research Center 1525 “Cardio-Immune interfaces” (funded by the German Research Foundation, grant number 453989101). R.S. is supported by a grant from the Canadian Institutes of Health Research (FDN-143299). This work was also supported by the Bavarian Ministry of Economic Affairs, Regional Development and Energy within the project “Single cell analysis in personalized medicine” at the Helmholtz-Institute for RNA-based Infection Research implemented in the Single-Cell Center Würzburg. The authors thank the Single-cell Center Würzburg for generating the RNA-seq data as well as the Core Unit FACS of the IZKF Würzburg for supporting this study. Figures were created with the help of BioRender.com.

## Author contributions

Conceptualization, G.C.R., D.A.; methodology, L.R., M.B., D.A.; formal Analysis, L.R., D.A., A.E.S., C.B., Z.M., R.S.; investigation, L.R., M.B., C.B., D.A., E.V.; writing – Original Draft, L.R., G.C.R.; visualization, L.R.; supervision, G.C.R.; funding Acquisition, G.C.R., D.A.

## Methods

### In silico analysis on the immune epitope database

The murine (*mus musculus*) protein sequence of myosin heavy chain alpha (MYHCA, product of the *Myh6* gene) was obtained from UniProt (protein accession number Q02566)^50^ and analyzed on the IEDB Analysis Resource for MHC-II binding prediction. Parameters were adjusted to predict binding of 15-mer peptides to murine MHC-II I-A^b^, found in C57Bl/6 mice. The MHC-II binding predictions were conducted in November 2017 using the IEDB analysis resource Consensus tool.^23^ The 10^th^ percentile was chosen as cutoff for selection of epitopes with high MHC-II binding affinity. For overlapping peptides sharing the same core sequence, those with the lowest adjusted rank value were selected for further analysis. Additional MHC-I binding predictions were conducted in January 2024 using the IEDB analysis resource SMM tool (Stabilized Matrix Method).^49^ In brief, the selected MYHCA sequence (YRILNPAAI) was queried for binding predictions on C57BL/6 MHC-I alleles H2-D^b^ and H2-K^b^. The predicted output was given in units of IC_50_ (nM).

### Mice

Wild type male C57BL/6J mice were purchased from Charles River (Jax Strain #:000664). Male and female B6.129S4-Ifng^tm3.1Lky^/J (C57BL/6 *Ifng*^YFP^, Jax strain #:017581) and C.Cg-Tg(DO11.10)10Dlo/J (BALB/c DO11.10, Jax strain #:003303) mice were imported from Jackson and bred at the Zentrum für Experimentelle Molekulare Medizin, at the University Hospital of Würzburg. The animals were maintained in individually ventilated cages under specific pathogen free (SPF) conditions with a 12-hour light/12-hour dark cycle and standard diet provided ad libitum. Experiments were conducted with 8- to 12-week-old mice. All procedures were conducted in accordance with the guidelines of the Federation for Laboratory Animal Science Associations (FELASA), the European animal welfare legislation, and were approved by the local authorities (*Regierung von Unterfranken*).

### In vitro stimulation of antigen-specific T cells from murine splenocytes

To assess potential T cell reactivity against defined cardiac peptides, splenocyte preparations were stimulated *in vitro* over 14 days with peptides of interest. In brief, animals were sacrificed by cervical dislocation and spleen and lymph nodes were harvested in ice-cold Hank’s balanced salt solution containing 0.5% (w/v) bovine serum albumin (HBSS/BSA) (Sigma Life Science, St. Louis, MO, USA). The lymphoid organs were then ground through a 30 μm filter mesh (Miltenyi Biotec, Bergisch Gladbach, Germany) and washed with HBSS/BSA (Sigma Life Science, St. Louis, MO, USA) to obtain a single cell suspension. Depending on downstream experiments, splenocytes were first subjected to magnetic activated cell sorting (refer to section ‘*Magnetic activated cell sorting’*) or used bulk for *in vitro* stimulation assays. After suspending cells in RPMI complete medium (10% FBS, 1% L-glutamine, 1% sodium pyruvate, 1% nonessential amino acids, 1% penicillin/streptomycin, 50 μM 2-mercaptoethanol (Gibco, ThermoFisher Scientific, Waltham, MA, USA) recombinant murine IL-2 (20 U/mL, PeproTech, Hamburg, Germany) and 1 μg/mL peptide (JPT Peptide Technologies, Berlin, Germany) were supplemented to the medium. 4*10^6^ cells were then cultured in 2 mL medium in 24-well plates at 37 °C and 5% CO_2_. As control, cells were cultured in the presence of IL-2 only. On days 4, 6, 8, and 11, 1 mL of culture medium was removed from each well and replaced with fresh RPMI complete medium containing IL-2 at 20 U/mL. On day 13, the cells were restimulated with the antigenic peptides of interest to induce a recall response. On day 14, the cells were finally harvested and subjected to flow cytometry or scRNA/TCR-seq analysis. The protocol used for the *in vitro* stimulation of cells was adapted from Chudley *et al.*.^51^

### Magnetic activated cell sorting

To characterize the MHC-restriction of the identified MYHCA epitope, CD4^+^ T cells were depleted from spleen and lymph node cell preparations. In brief, cells in suspension were stained with PE-conjugated anti-CD4 antibodies (clone GK1.5, ThermoFisher Scientific, Waltham, MA, USA), followed by labelling with magnetic anti-PE MicroBeads (Miltenyi Biotec, Bergisch Gladbach, Germany). Subsequently, the cell suspension was loaded on MACS-LS columns and magnetic activated CD4^+^ T cell depletion was performed in accordance with the manufacturer’s recommendations. CD4^+^ T cell depletion efficiency was routinely confirmed by flow cytometry analysis.

### Mouse IFN-γ/IL-2 Double-Color ELISpot Assay

Prior to ELISpot assay, splenocytes were *in vitro* stimulated with potential cardiac antigens as described elsewhere in the methods section and by Chudley *et al.*.^51^ On day 13 of stimulating culture, the cells were harvested and ELISpot assay was performed to detect IL-2 and IFN-γ-producing T cells in response to antigenic restimulation. The Murine IFN-γ/IL-2 Double-Color Enzymatic ELISpot Assay kit (CTL - Europe GmbH, Rutesheim, Germany) was used to quantify T cell responses according to the manufacturer’s instructions. In brief, the membrane in the 96-well-plates was activated by applying 70% ethanol, followed by thorough washing with PBS. Afterwards, the wells were precoated with IFN-γ/IL-2 capture antibodies, sealed and incubated at 4 °C overnight. After washing the plate with PBS, 50 µL of CTL-Test medium containing antigenic peptides (3 µg/mL) were added to each well. Positive control wells received medium with 1.6·10^6^ Dynabeads Mouse T cell activator CD3/CD28 beads/mL (Gibco, ThermoFisher Scientific, Waltham, MA, USA), while wells without peptides or Dynabeads served as negative controls. Cell suspensions (100 µL, 3·10^6^ cells/mL) were added to the ELISpot plate to reach a final peptide concentration of 1 µg/mL and a cell count of 3·10^5^ cells/well. The recall-assay plate was incubated at 37°C and 5% CO_2_ for 24 hours. Detection antibodies, enzyme conjugates, and substrates were then added for staining, with several washing steps in between. After air-drying the plate, the number of cytokine-producing cells was measured using an ImmunoSpot S5 Core ELISpot reader (CTL-Europe GmbH, Rutesheim, Germany). The count of cytokine producing cells was then corrected for background, introduced by the presence of IL-2 throughout culture.

### Flow cytometry

To stain cells for flow cytometry analyses, up to 10^6^ cells were transferred into U-bottom 96-well plates and washed with PBS (centrifugation at 300 g, 5’, 4 °C). The cells were subsequently stained with Zombie Aqua Fixable Viability Dye (1:1000, Biolegend, San Diego, CA, USA) or Fixable Viability Dye eFluor 780 (1:1000, ThermoFisher Scientific, Waltham, MA, USA) for 15 min at room temperature according to the manufactureŕs instructions. Following a washing step with FACS buffer (PBS containing 1% BSA, 0.1% sodium azide), the cells were incubated with fluorescently-labelled antibodies in the presence of anti-CD16/CD32 Fc blocking antibody (2.5 μg/mL, Biolegend, San Diego, CA, USA) for 30 min at 4 °C. The following antibody clones were commercially acquired from Biolegend (San Diego, CA, USA) and used at a final concentration of 2 μg/mL: anti-TCRβ (clone H57-597), anti-CD4 (clone RM4-5), anti-CD8α (clone 53-6.7) and anti-CD44 (clone IM7). After surface staining, the cells were washed with FACS buffer and resuspended thoroughly before proceeding to data acquisition. Flow cytometry analysis was conducted using an Attune NxT (Thermo Fisher Scientific, Waltham, MA, USA) instrument, and single-stain controls were used to compensate for spectral overlap. Data analysis was performed using FlowJo (FlowJo LLC Ashland, OR, USA, version 10.8.1).

### Cell sorting for single-cell RNA/TCR-sequencing

To perform 5’ single-cell RNA/TCR-sequencing on a desired cell population, cells were subjected to fluorescence-activated cell sorting (FACS) prior to sequencing. For this purpose, cells from *in vitro* stimulation cultures, were stained with viability dye and cell surface antibodies as described in section ‘*Flow cytometry’*. Sodium azide was omitted from the FACS buffer used for cell sorting purposes. Prior to FACS, cells were passed through a 35 μm filter mesh to remove residual cell clumps. Using a FACS Aria III instrument (BD, Heidelberg, Germany), cell populations originating from *in vitro* stimulation culture were sorted as follows: MYHCA_718-732_ stimulated live TCRβ^+^CD8^+^IFN-γ-YFP^+^CD44^high^ and IL-2 control live TCRβ^+^CD8^+^CD44^+^. The sorted cells were collected in tubes containing RPMI medium supplemented with 20% FBS. After sorting, IFN-γ-YFP^+^CD44^high^ (MYHCA_718-732_ stimulated) and CD44^+^ (IL-2 control) samples were labelled separately with anti-CD45/MHC-I TotalSeq^TM^-C C0301 and C0302 anti-mouse hashtag antibodies, respectively (1:150, 3.33 μg/mL, Biolegend, San Diego, CA, USA). Following two washing steps with FACS buffer and PBS 0.04% BSA, samples were resuspended in PBS containing 0.04% BSA and pooled. Finally, the multiplexed sample was loaded into the 10x Genomics Chromium pipeline for 5’ single-cell RNA/TCR-sequencing.

### Single-cell RNA/TCR-sequencing

Chromium™ Controller was used for partitioning single cells into nanoliter-scale Gel Bead-In-EMulsions (GEMs) and Chromium Next GEM Single Cell 5’ v1.1 kits for reverse transcription, cDNA amplification and library construction (10x Genomics, Pleasanton, CA, USA), following manufacturer’s instructions. A SimpliAmp Thermal Cycler was used for amplification and incubation steps (Applied Biosystems, Foster City, CA, USA). Libraries were quantified by a QubitTM 3.0 fluorometer (Thermo Fisher Scientific, Waltham, MA, USA) and quality was checked using a 2100 Bioanalyser with High Sensitivity DNA kit (Agilent, Santa Clara, CA, USA). Libraries were pooled and sequenced using the NovaSeq 6000 platform (S2 Cartridge, Illumina, San Diego, CA, USA) in paired-end mode to reach at least 70,000 reads per single-cell for gene expression and 7,000 reads for the T-cell receptor repertoire and hashtags. The Cell Ranger-7.0.0 Software suite (10x Genomics, Pleasanton, CA, USA) was used for sequence alignment, barcode processing and sample demultiplexing.

### In silico analysis of single cell sequencing data

Single-cell sequencing provided global transcriptome and TCR V(D)J sequence data.^52^ The gene count matrix obtained from Cell Ranger was analyzed using Seurat (version 4.0.4 ^53^). For sample demultiplexing, the detected cell barcodes in the gene expression data and hashtag oligo sequences were considered for each group. During quality control, cells with double hashtag signals or no hashtag detection were excluded from downstream analysis. Additional quality control parameters excluded cells yielding more than 60,000 unique molecular identifiers (UMI), more than 10% mitochondrial RNA or less than 200 expressed genes were also excluded. The cell transcriptome was then log-normalized, and the top 10,000 variable genes were selected for downstream steps. Next, principal component analysis (PCA) was performed on the scaled data. Twenty principal components obtained from PCA and a 0.4 resolution were then used to cluster cells. Ultimately, Uniform manifold approximation and projection for dimension reduction (UMAP) was applied to represent clusters in a nonlinear embedding. UMAP representation revealed six distinct cell clusters. The *FindallMarkers* function was then used to identify differentially expressed genes between clusters and to identify cluster biomarkers. Using the *combineExpression* function from the scRepertoire R package (version 1.2.2^54^), the productive, full-length V(D)J sequences for each T cell receptor were then added to the Seurat Object. After excluding cells with multiple or lacking α- or β-TCR chains, clones were called based on the CDR3 amino acid sequence. The Levenshtein distance was calculated for β-TCR sequences using the *adist()* function from the Utils R package, where a distance of 1 represents one amino acid insertion, deletion, or mutation. For calculation, TCR-sequences from the 20 most expanded clones from MYHCA_718-732_-expanded and IL-2-control condition were taken into account.

### GLIPH2

The Grouping of Lymphocyte Interactions by Paratope Hotspots (GLIPH2) algorithm was used to identify specificity groups in the TCR repertoire.^32,33^ By employing the *gliph_combined* function from the turboGliph R implementation (https://github.com/HetzDra/turboGliph), TCRβ CDR3 sequences from antigen-stimulated or control datasets that were at least three-times expanded were grouped based on the core motif of the CDR3 region. The terminal four amino acids on either end of the CDR3 sequences were excluded from analysis to avoid germline encoded V and J gene sequences and focus on the CDR3 core region, which has high contact probability with the antigenic peptide.^32^ A minimum length of three amino acids and a two-fold enrichment of the motifs in the sample set compared to a reference set, comprising 78,116 naïve murine TCRβ CDR3 sequences,^33^ was required for motif clustering. Additionally, a maximum probability score of 0.01 for the occurrence of a motif in an equal sized set of CDR3s from the control dataset was used as a cutoff.

### T cell activation reporter cell-line

BW5147.G.1.4 thymoma cells were purchased from ATCC (Manassas, VA, USA). Murine MSCV retroviral vectors for CD3-2A_pMI-LO (Addgene plasmid # 153418; http://n2t.net/addgene:153418; RRID:Addgene_153418)^38^ and 8xNFAT-ZsG-mCD8 (Addgene plasmid # 162747; http://n2t.net/addgene:162747; RRID:Addgene_162747)^39^ that were kindly deposited by Maki Nakayama were purchased from Addgene (Watertown, MA, USA) and purified from bacterial stocks. Retrovirus generation was performed using Platinum-E Retroviral Packaging Cell Line (Cell Biolabs, Inc., San Diego, USA). The endogenous TCR constant chains were knocked out using CRISPR/Cas9 technology. For this, BW-cells were transfected with TrueCut Cas9 protein v2 and TrueGuide Synthetic Guide RNA, specific for exon 1 of either TCR α or β constant chains, using the Neon NxT Electroporation System (all ThermoFisher Scientific, Waltham, MA, USA). After retroviral transduction, the surface expression of CD3 and CD8 was assessed by flow cytometry. Endogenous TCR KO was confirmed by lack of TCRβ expression in BW cells co-transfected with CD3 complex. Paired TCR α and β sequences were derived from single-cell sequencing and full TCR sequences were reconstructed from CDR3 regions, V genes and J genes using Stitchr.^40^ Genes for TCR α and β chains were separated using a T2A sequence. Full TCR gene blocks were synthesized and cloned into pMSCV-IRES-mThy1 vectors by VectorBuilder (Neu-Isenburg, Germany). After producing retrovirus, as described before, BW-TCR-KO-mCD3^+^-mCD8^+^-NFAT-ZsGreen reporter cells were transduced with TCRs of interest. TCR expression of reporter cells was confirmed by flow cytometry and TCRβ^+^ cells were sorted on a FACS Aria III instrument (BD, Heidelberg, Germany), before being used in downstream assays.

### Generation of murine bone marrow derived dendritic cells (BMDCs)

Syngeneic BMDCs were generated as previously described.^55^ Briefly, bone marrow precursor cells were obtained from flushed femurs and tibias of C57BL/6J mice and cultured at 3*10^6^ cells/ 10 ml in petri dishes in fully supplemented RPMI and rmGM-CSF (200 U/ml; Peprotech, Hamburg, Germany). Medium was exchanged on days 3 and 6 and the cells were cryopreserved on day 7 for further use.^56^

### Co-culture of BW-NFAT-ZsGreen cells with BMDCs

BW-NFAT-ZsGreen reporter cell lines with reconstructed TCRs were co-cultured with BMDCs loaded with antigen to probe for TCR antigen specificity. BMDCs were thawed into 200 U/ml GM-CSF-containing RPMI complete medium (200 U/ml; Peprotech, Hamburg,Germany) in 10 cm petri-dishes one day in advance. For co-culture, cells were plated at a ratio of 4 reporter T cells to 1 BMDC in 96 well plates in the presence of MYHCA_724-732_ (YRILNPAAI) and OVA_257-264_ (SIINFEKL) peptides (JPT Peptide Technologies, Berlin, Germany). After 24 h or 72 h, cells were stained for viability, CD8α and TCRβ and analyzed by flow cytometry for NFAT-ZsGreen reporter activity.

### Statistics

The graphs show the individual distribution of *n* datapoints alongside with mean ± S.E.M. Statistical analysis was done using unpaired t test with correction for multiple comparisons by Holm-Sidak method, One-way ANOVA with Dunnett’s multiple comparisons test, or Two-way ANOVA followed by Tukeýs multiple comparisons test. Differences were considered statistically significant for P < 0.05. *, ** and *** indicate P < 0.05, P < 0.01 and P < 0.001 respectively. Statistical analysis was conducted using Graph Pad Prism (v. 10.2.1).

## Supplemental information

Document S1: Figures S1, S2, S3 and Tables S1, S2

## Resource availability

### Corresponding authors

Further information and requests for resources and reagents should be directed to and will be fulfilled by the corresponding authors, Prof. Dr. Gustavo Campos Ramos and Dr. DiyaaElDin Ashour.

### Materials availability

This study did not generate new unique reagents.

### Data and code availability

The transcriptomic data acquired in this study will be made available on NCBÍs Gene Expression Omnibus database after the peer-reviewed version is published. Any additional information required to reanalyze the data reported in this paper is available from the corresponding authors upon request.

