## Supplemental Figures 1-3 for "Identification of a novel myosin antigen activating CD8^+^ T cells in C57BL/6 mice"

[*] **Corresponding authors**

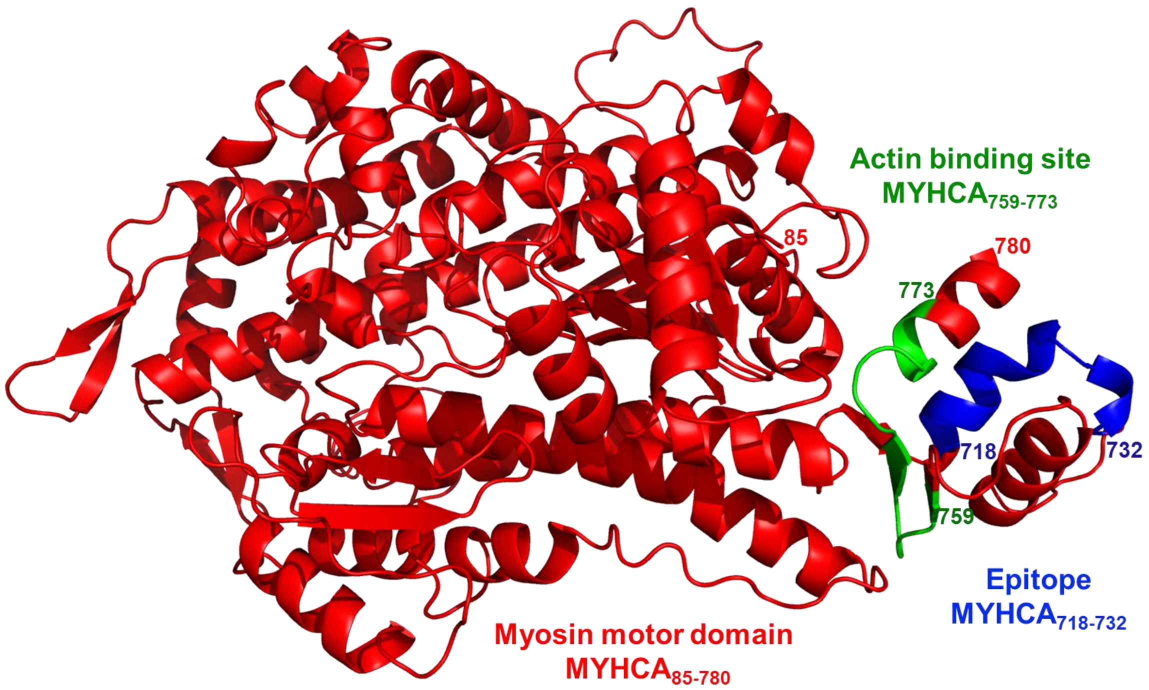


**Figure S1: MYHCA protein structure.** Three-dimensional representation of the motor domain of myosin S1 fragment is shown in red. Identified novel epitope ($MYHCA_{718\text{-}732}$) and the adjacent actin-binding site of myosin ($MYHCA_{759\text{-}773}$) are shown in blue and green, respectively. The structure was predicted by AlphaFold (AF-Q02566-F1) using mouse Myosin heavy chain 6 (MYHCA) sequence from UniProt (UniProtKB-Q02566) and visualized by PyMOL.

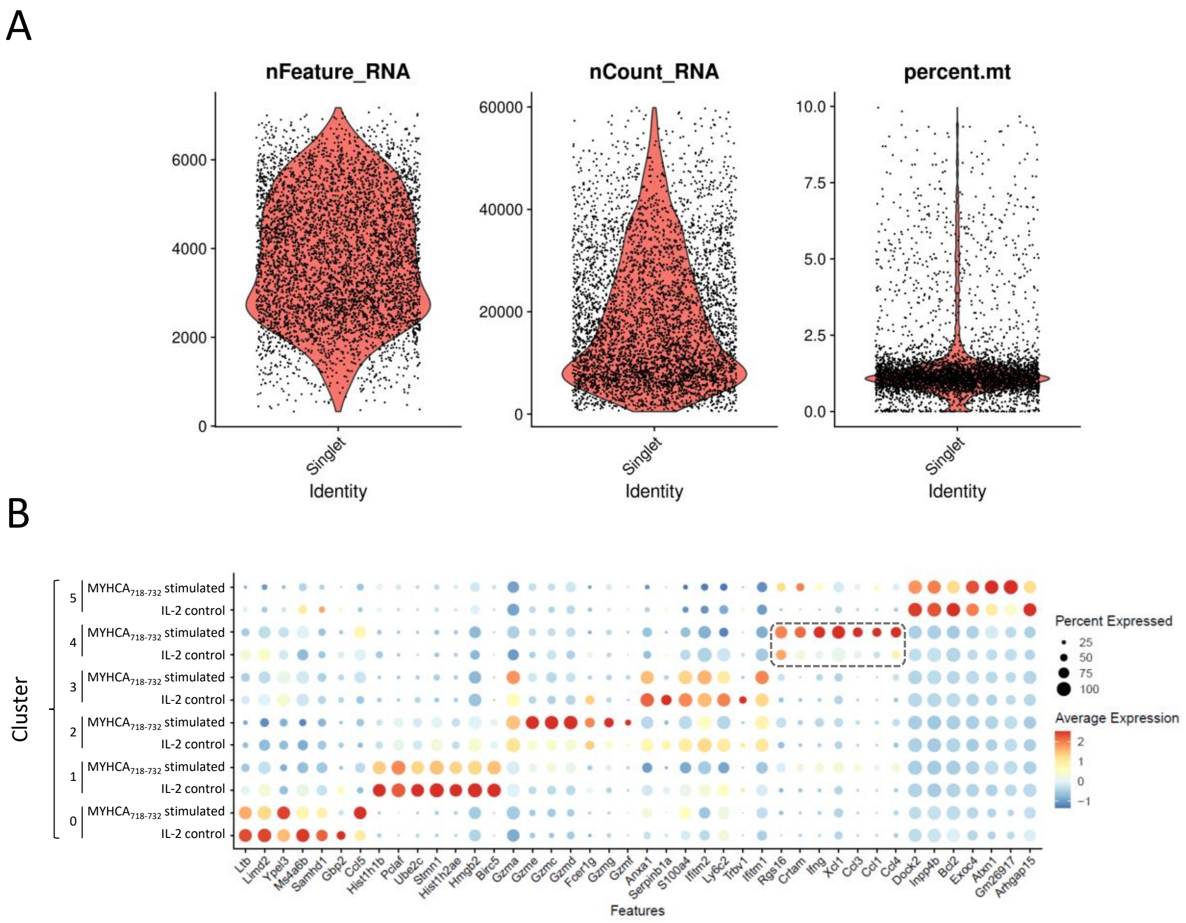


**Figure S2: scRNA/TCR-seq quality control and differential gene expression. A:** Quality control of single-cell RNA-seq data. Violin plots display the number of expressed genes (nFeature_RNA), the RNA count (nCount_RNA) and the percentage of mitochondrial genes (percent.mt) per cell in single-hashtag-cells from the single-cell object. Quality control excluded cells yielding less than 200 expressed genes, more than 60,000 unique molecular identifiers (UMI) or more than 10% mitochondrial RNA. **B:** Differential gene expression. Dot plot presenting the seven most differentially expressed genes among all identified clusters, split by IL-2 control and $MYHCA_{718-732}$-stimulated condition. Gene expression corresponding to cluster 4 is highlighted.

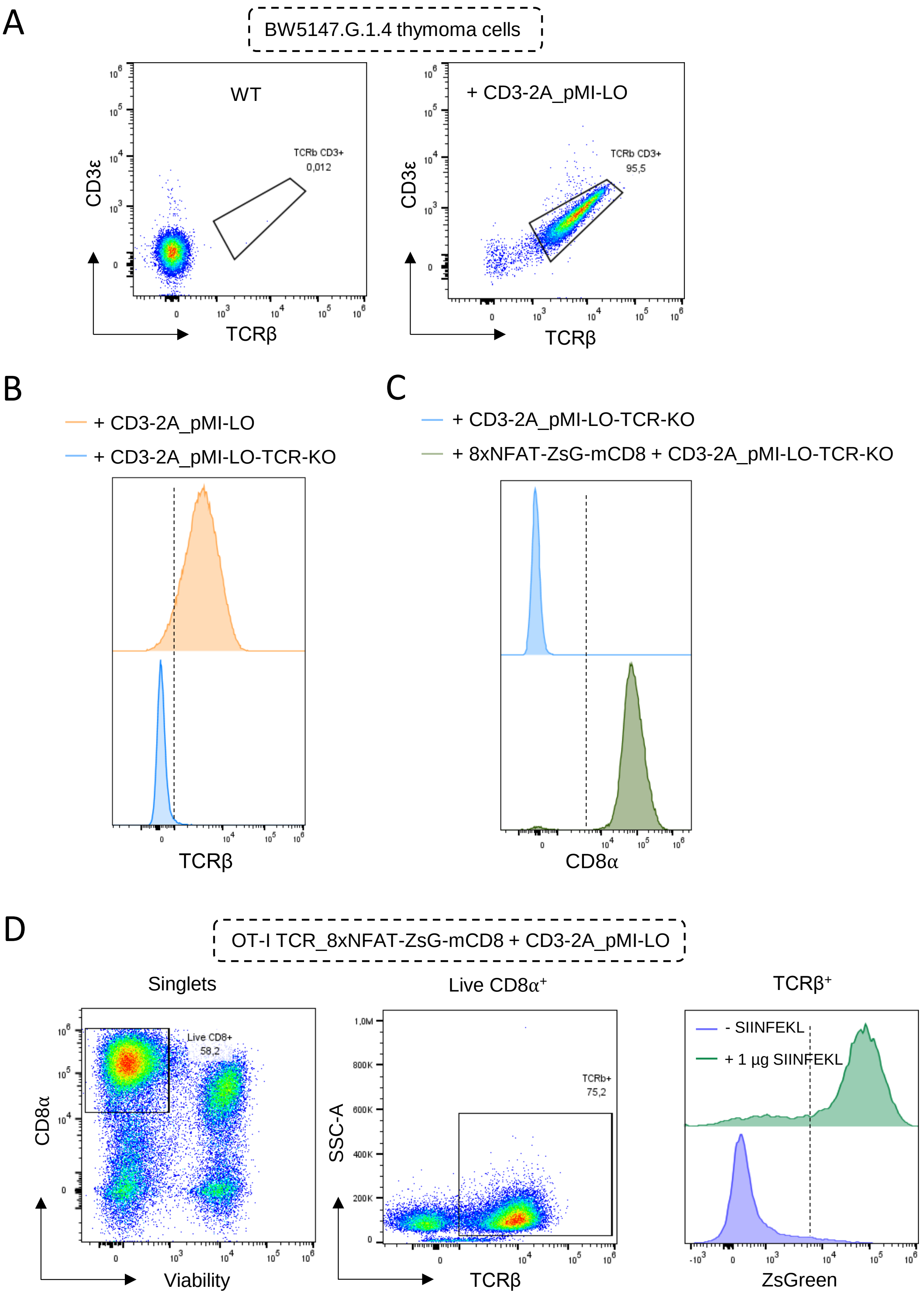
A
BW5147.G.1.4 thymoma cells
WT
TCRb CD3+
0,012
CD3ε
TCRβ
+ CD3-2A_pMI-LO
TCRb CD3+
95,5
CD3ε
TCRβ
B
+ CD3-2A_pMI-LO
+ CD3-2A_pMI-LO-TCR-KO
TCRβ
C
+ CD3-2A_pMI-LO-TCR-KO
+ 8xNFAT-ZsG-mCD8 + CD3-2A_pMI-LO-TCR-KO
CD8α
D
OT-I TCR_8xNFAT-ZsG-mCD8 + CD3-2A_pMI-LO
Singlets
Live CD8+
58,2
CD8α
Viability
Live CD8α+
TCRb+
75,2
SSC-A
TCRβ
TCRβ+
- SIINFEKL
+ 1 µg SIINFEKL
ZsGreen

**Figure S3: Generation of TCR-KO-$CD3^+CD8^+$ BW5147.G.1.4 NFAT-ZsGreen reporter line.** **A:** Transduction of CD3-2A_pMI-LO construct into BW5147.G.1.4 thymoma cells. The endogenous TCR of BW5147.G.1.4 thymoma cells is re-expressed on the surface. **B:** Histogram of TCR-$\beta$ before (orange) and after (blue) CRISPR/Cas9 mediated excision of TCR $\alpha$ and $\beta$ constant chains from BW5147.G.1.4 + CD3-2A_pMI-LO line. **C:** Transduction of 8xNFAT-ZsG-mCD8 construct into TCR-KO BW5147.G.1.4 + CD3-2A_pMI-LO line. **D:** Gating strategy for ZsGreen signal in OT-I TCR_8xNFAT-ZsG-mCD8 + CD3-2A_pMI-LO line after co-culture with unloaded or with or without 1 µg/mL $OVA_{257-264}$-loaded BMDCs.

### Supplemental tables

**Table S1:** Amino acid-sequences of MYHCA-derived candidate antigens revealed by *in silico* prediction (A) and MYHCA$_{718-732}$ 9-mer peptides (B).

| | **Peptide position** | **Amino acid sequence** |
|---|---|---|
| A | MYHCA$_{111-125}$ | AWMIYTYSGLFCVTV |
| | MYHCA$_{126-140}$ | NPYKWLPVYNAEVVA |
| | MYHCA$_{189-203}$ | KRVIQYFASIAAIGD |
| | MYHCA$_{248-262}$ | FIRIHFGATGKLASA |
| | MYHCA$_{307-321}$ | NPYDYAFVSQGEVSV |
| | MYHCA$_{347-361}$ | KAGVYKLTGAIMHYG |
| | MYHCA$_{418-432}$ | VQQVYYSIGALAKSV |
| | MYHCA$_{434-448}$ | EKMFNWMVTRINATL |
| | MYHCA$_{718-732}$ | GDFRQRYRILNPAAI |
| | MYHCA$_{720-734}$ | FRQRYRILNPAAIPE |
| B | MYHCA$_{718-726}$ | GDFRQRYRI |
| | MYHCA$_{719-727}$ | DFRQRYRIL |
| | MYHCA$_{720-728}$ | FRQRYRILN |
| | MYHCA$_{721-729}$ | RQRYRILNP |
| | MYHCA$_{722-730}$ | QRYRILNPA |
| | MYHCA$_{723-731}$ | RYRILNPAA |
| | MYHCA$_{724-732}$ | YRILNPAAI |

**Table S2:** TCR CDR3 sequences from expanded MYHCA$_{718\text{-}732}$-stimulated clones (clone frequency ≥ 1% in stimulated subset)

| Clone | CDR3 amino acid sequence ($\alpha$_$\beta$) |
|---|---|
| 1 | CSASRDYSNNRLTL_CASSIGGQDTQYF |
| 2 | CALENSGGSNAKLTF_CASSLVTGGVEQYF |
| 3 | CAMGITGNTGKLIF_CASSSIYEQYF |
| 4 | CATVSNTGYQNFYF_CASSLDWGDYAEQFF |
| 5 | CAVISSGSWQLIF_CASTRDWGYEQYF |
| 6 | CAVASSGSWQLIF_CASNRDWGYEQYF |
| 7 | CALSDRTEGADRLTF_CASSLDWGSNYAEQFF |
| 8 | CAVRGNMGYKLTF_CASSPDWGVSYEQYF |
| 9 | CALGEGGADRLTF_CASSLEWGGSYEQYF |
| 10 | CALSDDTNAYKVIF_CASSLEPSGNTLYF |
| 11 | CVLSANSNNRIFF_CASSELSYEQYF |
| 12 | CALGALPSFSKLVF_CASSLVLSQNTLYF |
| 13 | CALSGGGSALGRLHF_CASSLGQHTEVFF |
| 14 | CALEPGTGYQNFYF_CAWSLQNSYNSPLYF |
| 15 | CAVKNNTNTGKLTF_CASSPDWGVSYEQYF |
| 16 | CAVSGASSGQKLVF_CASSLDWGSSYEQYF |
| 17 | CAVNSGGNYKPTF_CASSLAQGGTGQLYF |
| 18 | CALISNNRIFF_CASSNSYEQYF |
| 19 | CAVSIRGRALIF_CASSESNWATGQLYF |
| 20 | CALGDRSSNTNKVVF_CASRDNYAEQFF |
